# Relative effects of edaphic conditions and climate on palm communities in the Central Andes

**DOI:** 10.1101/2021.03.08.434423

**Authors:** Fabiola Montoya, Moraes R. Mónica, Alfredo F. Fuentes, Leslie Cayola, Ana Antezana, Tatiana Miranda, Esther Mosqueira-Meneses, M. Isabel Loza, J. Sebastián Tello

## Abstract

Palms (family Arecaceae) are conspicuous and structural elements in forests ecosystems of tropical regions and mountain forests in South America. Additionally, many species of palms are culturally and economically important to human populations. Because of their ecological and ethnobotanical significance, understanding the drivers of palm distribution and diversity is critical. However, most past research has focused in tropical lowland palm communities, while our understanding of montane tropical palm ecology and biogeography is comparatively lacking. We investigate the environmental factors that influence patterns of richness, composition, and abundance of palms in the Central Andes. In particular, we are interested in the relative effects that soil edaphic conditions and climate have on palm community structure. For these analyses, we used a network of 88 forest plots distributed along a broad elevational gradient (1,000 – 3,200 meters), which are part of the Madidi Project in north-western Bolivia. We carried out palm community-level analysis, as well as species-specific analyses for each of the 16 most common species. We found that soils and climate contribute differentially to shaping Andean palm diversity and distributions. We found that soils explain more variation in species composition (14.4%) than climate (3.45%), but that climate explains more variation in species richness (13%) than soils (6.1%). Moreover, species-specific analyses reveal that there is great variation in how different common species respond to their abiotic environment. Our results contribute to understanding the drivers of biodiversity for a highly important group of plants in one of the most important hotspots for biodiversity.

**RESUMEN:** En el neotrópico, las palmeras (Arecaceae) son un grupo diverso y abundante de plantas que constituyen elementos estructurales en bosques tropicales tanto de tierras bajas como de montaña. Además, muchas especies de palmeras son culturalmente y economicamente importantes para muchas poblaciones humanas. Debido a su importancia ecológica y etnobotánica, entender los mecanismos que controlan la diversidad y la distribución de las palmeras es extremadamente importante. Sin embargo, la mayoría de la investigación hasta el momento se ha enfocado en comunidades de palmeras de tierras bajas, mientras que la ecología y biogeografía de las palmeras de montañas es relativamente poco entendida. En este estudio, nosotros investigamos los factores ambientales que influencian la riqueza, composición y abundancia de palmeras en los Andes Centrales. En particular, estamos interesados en entender los efectos relativos de las condiciones edáficas del suelo y el clima en la estructura de comunidades de palmeras. Para nuestros análisis, usamos una red de 88 parcelas de árboles distribuidas a lo largo de un gradiente elevacional (1,000 – 3,200 metros), la misma que es parte del Proyecto Madidi en Bolivia. Encontramos que el suelo y el clima tienen efectos diferentes. El suelo explica más variación en la composición de especies (14.4%) que el clima (3.45%), pero el clima explica más variación en la riqueza de especies (13%) que los suelos (6.1%). Además, análisis independientes para las 16 especies más comunes demuestran gran variación en como cada especie responde a las condiciones ambientales. Nuestros resultados contribuyen a entender los factores que controlan la diversidad de un grupo importante de plantas en una de las regiones más diversas del planeta.

## 1. INTRODUCTION

Palms (family Arecaceae) are a charismatic and diverse group of plants, which play critical ecological roles in tropical ecosystems, and provide many important natural services to human populations. Many palms are important elements of tropical and montane forests across South America, contributing significantly to the structure and diversity of uppermost canopy, as well as of the mid- and understorey (Svenning 2001, Henderson 2002, Garibaldi & Turner 2004, Eiserhardt *et al*. 2011, Balslev *et al*. 2012). In these ecosystems, palm species provide food and shelter, and their local abundance can make them a keystone resource for populations of vertebrate and invertebrates animals (palm fruits are edible, either for their soft mesocarp or for their endosperm (Zona & Henderson 1989, Muñoz *et al*. 2019). Palms also have many important ethnobotanical uses for human societies across the tropics. For example, people use stems as poles in house construction and leaves for thatching roofs. Leaves also provide fibers for weaving baskets, fishing nets and other tools (Paniagua-Zambrana *et al*. n.d., Garibaldi & Turner 2004). Other palms are used in ceremonial or religious activities and other activities of cultural importance (Paniagua-Zambrana *et al*. n.d.). Indeed, palms species provide fuel, fabrics and medicine for many rural and urban communities (Paniagua-Zambrana *et al*. n.d., Henderson 2002, Eiserhardt *et al*. 2011, Montoya & Moraes R. 2014, Moraes R. 2015). For all these reasons, natural populations of palms are exploited, and their environments are threatened. Thus, understanding the drivers of the amazing geographic variation in palm richness, composition and abundance is fundamental to conservation strategies focused on their preservation.

Multiple evolutionary and ecological forces have been proposed to explain variation in the diversity and composition of natural communities. These processes range in scale from geological changes such as mountain uplift that creates new habitats and physical barriers for dispersal (Pintaud *et al*. 2008) to local interactions such as apparent competition mediated by shared enemies (Pintaud *et al*. 2008, Eiserhardt *et al*. 2013). However, one of the most important factors that shape the distribution and abundance of species, and that often mediate the effects of biogeographic history and species interactions, is the abiotic environment (Condit *et al*. 2013). In this way, understanding the effects abiotic conditions on community diversity and composition is often a fundamental first step needed before deeper insights can be obtained. This is the main objective of our study.

Past research has suggested that climate and soils are important drivers of plant communities. For example, (Eiserhardt *et al*. 2013)showed that habitat types, characterized by inundation regime, play an important role in the spatial turnover of species in Amazonian rainforests (Sesnie *et al*. 2009, Eiserhardt *et al*. 2013). In particular palms are well known as a tropical clade of species that strongly prefer wet and warm conditions (Eiserhardt *et al*. 2011). Thus, we expect that precipitation and temperature will be important drivers of palm abundance across elevational gradients. Indeed, Sesnie *et al*. (2009) found that precipitation affect the palm abundance in north-eastern Costa Rica, while Costa *et al*. (2009) suggested that water drainage patterns seem to be a major factor controlling patterns of palms distribution along topographic gradients. Precipitation might be particularly important in montane regions because high topographic heterogeneity can cause dramatic variation in precipitation from place to place (Eiserhardt *et al*. 2011).

Plant distributions also often track soil conditions, particularly soil nutrient availability. For example, Figueiredo *et al*. (2018) found that soils are a more important predictor of plant species ranges in Amazonia than climate. Their study suggests that species distribution are limited by edaphic factors that reduce species’ abilities to track suitable climate conditions. In a different study, (Condit *et al*. 2013) found that soil phosphorus was the strongest factor affecting the distribution of more than half of the species of tropical vegetation in central Panama. The finding that many species have associations with either high or low phosphorus reveals an important role for this nutrient in limiting tropical tree distributions. The distribution of palm species has also been associated with soil conditions like texture and nutrients. For example, Phillips *et al*. (2003) and Tuomisto *et al*. (2003) found an important role of soil properties in turnover of species composition within *terra firme* forests.

Climate and edaphic conditions can sometimes interact. Indeed, changes in temperature, precipitation or topography (such as slope and aspect) can play a big role in shaping variation in soil conditions and nutrient availability. For example, temperature controls microbial activity, which in turn reduce the decomposition rate (Lloyd & Taylor 1994), affecting the total amount of nutrients available and organic carbon stocks (Garibaldi & Turner 2004, Girardin *et al*. 2010, Figueiredo *et al*. 2018). For all these reasons, to understand the effects of soils we need to simultaneously understand the effects of climate on plant distributions.

Many studies have investigated the effect of climate and edaphic conditions on palm distribution in lowlands (e.g., the Amazon) or premontane forest (Jones *et al*. 2008, 2013, Pintaud *et al*. 2008, Eiserhardt *et al*. 2011, 2013, Kristiansen *et al*. 2012, Condit *et al*. 2013, Prada *et al*. 2017, Schlindwein *et al*. 2017). In contrast, there is a paucity of studies that simultaneously consider the effects of edaphic and climate conditions on palms communities in the Andes. This is so, despite a high diversity of palms in montane regions. Indeed, the tropical Andes above 1,000 m host 121 species of palms and 24 genera (Pintaud *et al*. 2008, Arias & Stauffer 2012). Just as in lowland forests, palms can constitute an important component of the structure and ecology of montane forest (Moraes *et al*. 1995, Borchsenius & Moraes 2006).

In this study, we used 88 forest plots distributed along a 2,200 m elevational gradient to assess the effects of environmental conditions (climate and soils) on Andean palm species communities. Specifically, we address three interrelated question: (1) Is there an elevational gradient in the diversity, abundance and species composition of Andean palm communities? We expect that palm communities will change across the elevation gradient in response to diverging environmental conditions. (2) Do environmental conditions, particularly climate and soils, explain variation in the diversity, abundance and composition of Andean palm communities?

This question explores the relative contribution of the environmental conditions on the structure of communities. (3) Do climatic and soil conditions explain spatial variation in abundance of the most common species of Andean palms? Here, we study how individual palm species respond to their abiotic environment to understand the factors that have species-specific effects on abundance patterns. We found that soils explain more variation in species composition (14.4%) than climate (3.45%), and climate explains more variation species richness (13%) than soils (6.1%). Species-specific analyses reveal that there is great variation in how different common species respond to their abiotic environment. Our results contribute to understanding the drivers of biodiversity for a highly important group of plants in one of the most important hotspots for biodiversity.

## 2. METHODS

### 1. Species composition and abundance data

In this study, we used data from the Madidi Project (https://madidiproject.weebly.com/), which includes a large network of nearly 500 forest plots distributed along an elevational gradient in the Amazon and Andes of northwestern Bolivia. Because we were interested in studying Andean palm communities, our analyses used only 88 plots located above 1,000 m in elevation that contained at least one palm individual. These plots are 0.1-ha in area (50 x 20 m), are typically located in mature forest at least 500 m apart from one another and cover a broad range of environmental conditions ranging in elevation from 1,000 to 3,200 m (Figure 1; (Tello *et al*. 2015)). Mean annual temperature varies from 14.6 to 21.6°C, while total annual precipitation ranges from 652.8 to 2,882 mm (climate data from WorldClim 2.0; (Fick & Hijmans 2017)). The soils in the region can vary from moderately deep to very deep, and can be alkaline, silty-clay-loam, silty-clay, loamy-clayey and clayey, in addition to frequent gravel and deep stones (Fuentes 2005). Within our forest plots, all palm individuals with a diameter at breast height (DBH) equal or greater than 2.5 cm were measured and taxonomically identified. Herbarium specimens to document each species at each site are deposited mainly at the Herbario Nacional de Bolivia and the Missouri Botanical Garden. All palms were identified to species level. In total, we recorded 3,148 individuals belonging to 16 species. The data used in our analyses correspond to version 2.2 of the Madidi Project Dataset, which is deposited in Zenodo (DOI 10.5281/zenodo.4280178) (Tello *et al*. 2018).

**FIGURE 1.**
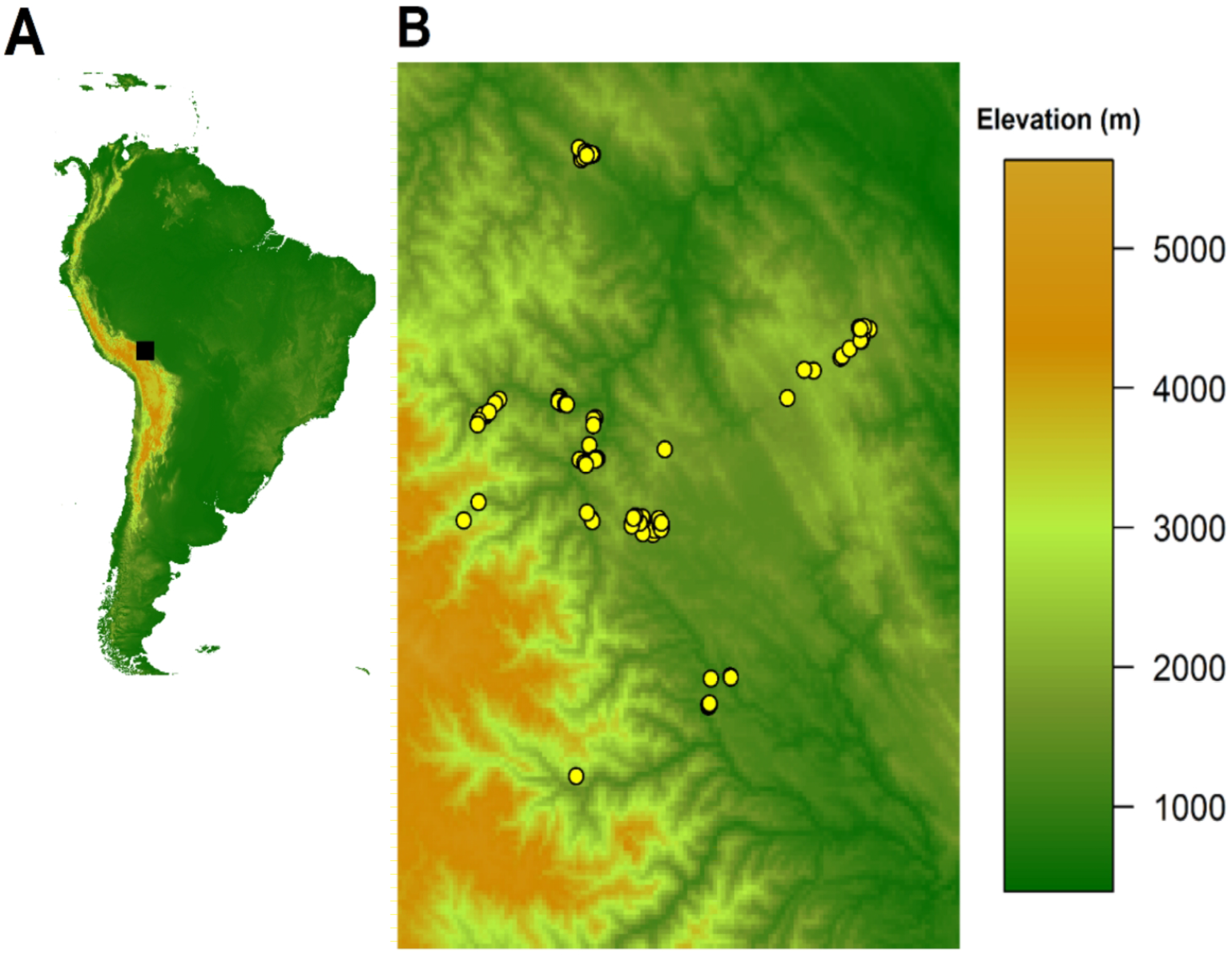
Location of the study region and forest plots used in this study. (A) The Madidi Region is located in the areas in and around the Madidi National Park in northwestern Bolivia (B) The 88 temporary plots (yellow dots) with palm abundance data are distributed along an elevational gradient from 1,000 to 2,500 m.

From these data, we constructed a community matrix that contains 16 palm species (columns) distributed across 88 plots (rows). This matrix is filled with the abundance of each species at each site. For each plot, we calculated the number of palm species (richness) and the total number of individuals (total abundance). We then transformed the community matrix using the Hellinger method described by Legendre & Gallagher (2001) and implemented by the function “*decostand”* in R package “*vegan*” (Oksanen *et al*. 2016). This transformation reduces the weight of rare species on the estimation of dissimilarities between plots and is recommended to diminish bias in ordination methods (Legendre & Gallagher 2001). Moreover, when used in combination with methods based on Euclidean distances, such as principal component analysis or redundancy analysis, this transformation avoids the double-zero problem (Legendre & Gallagher 2001).

### 2. Climate predictors

We used the geographic coordinates of each plot to extract values for 19 bioclimatic variables from WorldClim Version 2.0 (Fick & Hijmans 2017). These variables reflect the annual average and temporal variability in temperature and precipitation. The WorldClim dataset is generated by an interpolation of long-term monthly temperature and precipitation data (mostly for the 1970-2000 period) from weather stations in a large number of global, regional, national and local sources. The interpolation uses satellite-derived weather data and other covariables to create global climate surfaces (Fick & Hijmans 2017).

To reduce the dimensionality of the climate dataset, we employed a principal component analysis (PCA) on centered and standardized variables. The first three PC axes accounted for 93% of variation in climate (Figure S1A), and only these were retained for further analyses. The first PC axis (hereafter Clim1) is dominated by positive loadings for mean and extreme temperature and by negative loadings of temperature variability (Figure S1B). Thus, Clim1 reflects variation from colder sites with large temperature variation to warm sites that change little in temperature through the year. The second PC axis (Clim2) reflects a variation in precipitation from dry to wet sites (Figure S1C). Finally, the third PC axis (Clim3) represents a gradient from sites with low precipitation seasonality and high precipitation in the driest part of the year to sites with high seasonality in precipitation and less rain during the driest part of the year (Figure S1D, Table S1).

### 3. Soil predictors

Soil characteristics were measured in samples collected during the vegetation census of each plot. Within each plot, three sub-samples were taken at randomly selected locations between trees. The samples were taken from a 50 × 50 cm square, free of leaf litter and stones. Each sub-sample was taken from a depth of 0 to 30 cm below the litter layer, and then combined and mixed into a full sample of 1 kg. All soil samples were analyzed by the Laboratorio de Calidad Ambiental of the Instituto de Ecología at the Universidad Mayor de San Andrés in La Paz, Bolivia. Soil conditions were characterized by 11 variables (including 6 nutrients). Total nitrogen (N) was measured using the semi-micro Kjeldahl method, while phosphorus (P), sodium (Na), potassium (K), calcium (Ca) and magnesium (Mg) using the 1M ammonium acetate solution method. We also measured pH and electro-conductivity (EC) with a potentiometer, cationic exchangeable capacity (CEC) was measured using the summation of total bases. Finally, silt and sand percentages were estimated with the hydrometer method. Clay percentage was excluded because of its perfect multicollinearity with silt and sand (Text S1).

To reduce dimensionality, we log-transformed all soil variables and conducted a PCA on centered and standardized variables. For all subsequent analyses, we used the first seven principal components, which accounted for 92% of the variation in edaphic conditions (Figure S2A). We determined that the first PC axis (Soils1) represents a gradient from low to high concentrations in most micronutrients (P, Mg, Ca and K; Figure S2B). The second PC axis (Soils2) represents a gradient from silty soils to sandy soils with high Na concentrations. Soils3 is a gradient from acidic soils to neutral soils with high concentration in N and high electric-conductivity (Figure S2D). Soils4 characterizes a gradient from high to low Na concentrations and cationic exchangeable capacities (Figure S2E). Soils5 represents a gradient from high percentages of silt and sand (i.e. low clay) and high concentration of K to soils with high electro-conductivity and high percentages of clay (Figure S2F). Soils6 characterizes sites with high P concentration to sandy sites with high cationic exchangeable capacities (Figure S2G). Finally, Soils7 represents a gradient from low to high concentrations of P and low to high cationic exchangeable capacity (Figure S2H). For more details, see Tables S2 and S3.

### 4. Statistical analyses

In order to address our first question, we regressed palm species richness and total abundance separately against elevation. For these analyses, we used linear and quadratic ordinary least square models (OLS) and compared them using an ANOVA. If the quadratic model was significantly better, it was retained and used for interpretation (Table 1). In addition, we conducted a redundancy analysis (RDA) where the Hellinger-transformed matrix of composition was the response, and the linear and quadratic terms of elevation were predictors. For these analyses, we used the function “*rda*” in the R Package “*vegan*” (Oksanen *et al*. 2016).

**Table 1.**
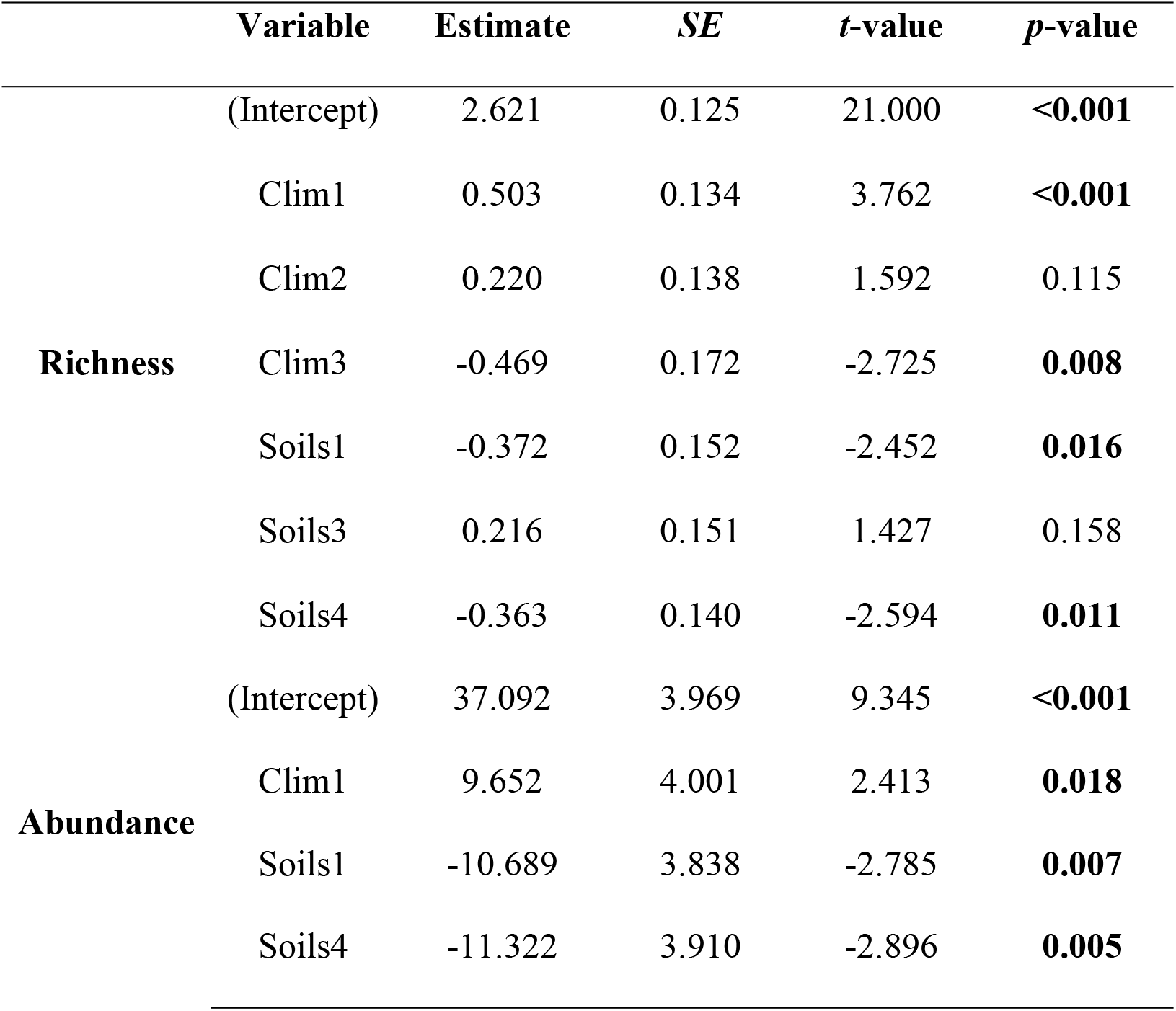
Most important climate and soil predictors for richness and abundance. The table shows result of spatial GLS regressions with the predictors that were selected based on a search for a minimum most adequate model using AIC (see methods). These analyses were conducted separately for palm richness and total palm abundance. Predictors includes principal components derived from climate and soil data (see Methods). For each predictor in each model, a coefficient estimate, standard error (*SE*), *t-*value and *p*-value are reported. Bold font highlights statistically significant predictors.

For our second question, we conducted separate analyses for (1) species richness, (2) total abundance and (3) the Hellinger-transformed matrix of species composition. First, we conducted a series of variation partitioning analyses (function “*varpart*” in R package “*vegan”*). These analyses partition the variation in the response into fractions explained by different sets of predictors. In our case, we partitioned the variation (in richness, abundance or composition) into fractions explained (1) only by soils, (2) only by climate, (3) simultaneously by both soils and climate (i.e shared variation) and (4) the variation that remains unexplained. In each analysis, we used all three climate principal components and all seven soil principal components as predictors.

Additionally, we constructed minimum adequate models (MAMs) for each response. For species richness and total abundance, we used spatial generalized least square models (GLS; details below), while for community composition we used a redundancy analysis. In both cases, models started with a full complement of predictors (all climate and soil principal components) and the model selection was based on a stepwise procedure using the Akaike information criterion (AIC). For model selection, we used the function “*stepAIC*” in the R package “*MASS”* (Venables & Ripley 2002). The final model (the MAM) contains the combination of predictors that most reduce the AIC value of the model, and that are likely the most important factors responsible for variation in the response variable. All predictors were centered and standardized to make regression coefficients comparable.

As mentioned above, for richness and total abundance, we used generalized least-squared (GLS) models. These models used geographic coordinates of each plot to account for spatial autocorrelation among sites. We used a Gaussian correlation structure in all analyses because it was the best fit for our models (functions “*gls”* and “*corGaus”* in the R package “*nlme*”; (Pinheiro *et al*. 2020)). After producing the minimum adequate model (MAM) by AIC selection, we compared this model against a null model. This allowed us to obtain a model-wide p-value for the MAM via ANOVA. The null model was another GLS regression that only contained an intercept term and the same spatial autocorrelation structure estimated for the MAM. As estimates of relative fit, we calculated the difference in AIC between the MAM and the null model, as well as a pseudo-R^2^ using the function “*rsquared*” in the R package “*piecewiseSEM”* (Lefcheck 2016).

Unfortunately, there is no GLS equivalent when using a multivariate response. Thus, to study the climatic and edaphic determinants of species composition we used a redundancy analysis (RDA). Here, the Hellinger-transformed matrix of community composition was the response and climatic and edaphic PC axes were predictors. For the redundancy analysis, an AIC cannot be computed, so the variable selection is based on a metric that mimics AIC but is based on the residual sums of squares (function “*deviance*.*cca*” in R package “*vegan*”). After variable selection, an adjusted *R*^*2*^ and *P*-values were obtained directly from model output.

For our third question, we related the abundance of each species separately with climate and soil predictors. Not all palm species had sufficient data for these analyses; thus, we only used the seven species that occur in at least 15 plots: *Aiphanes horrida* (26 plots), *Chamaedorea angustisecta* (16), *Dictyocaryum lamarckianum* (41), *Euterpe precatoria* (52), *Geonoma orbignyana* (29), *Iriartea deltoidea* (19) and *Socratea exorrhiza* (28). For all analyses, we retained only the plots where the focal species was present (i.e., eliminated all zeroes). Additionally, because of the reduced sample size, we only considered the first three principal components of climate and the first four principal components of soils. Including more predictors would likely lead to overfitting each model. Just like for the analyses on richness and total abundance, analyses for each species involved (1) a variation partitioning procedure using all predictors and (2) the construction of a minimum adequate model using AIC variable selection and spatial GLS models. The details of these methods are the same as those described for question two.

## 3. RESULTS

Is there an elevational gradient in the diversity, abundance and species composition of palm communities? Species richness showed a humped-shaped relationship with elevation (i.e. quadratic regression better than lineal regression; Figure 2A; *F* = 7.04; *P* = 0.002). Richness reached its maximum between 1,200 – 1,600 m and decreased to only one species above 2,200m. In total, elevation explained 11.4% of variation in species richness. In contrast, abundance was not significantly related to elevation (neither the quadratic nor linear regressions were significant; Figure 2B; *P* > 0.05; *R*^*2*^ = 0.014). Finally, the RDA showed that elevation (linear and quadratic terms) explains a small proportion of variation in palm species composition among plots (6.3%; Figure 2C).

**FIGURE 2.**
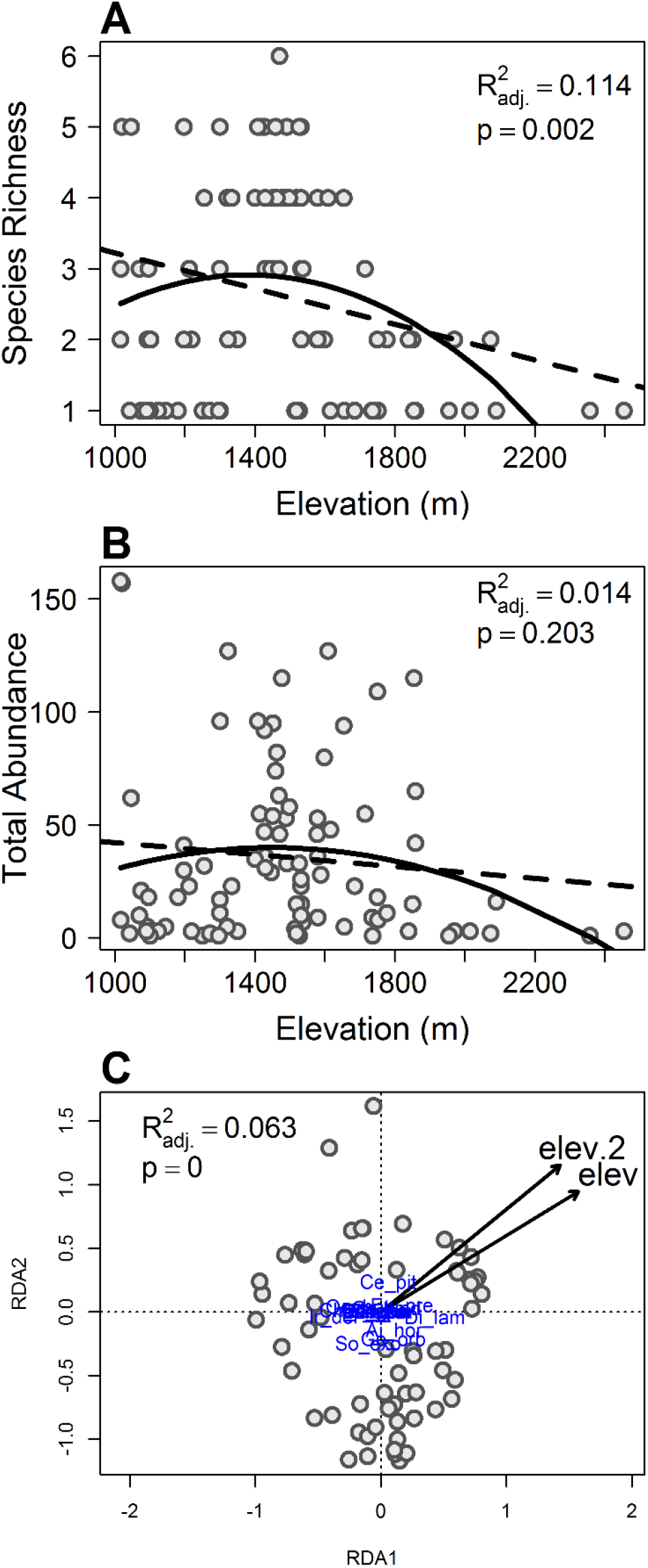
Variation in palm species richness, total abundance and composition in relation to elevation. (A) Palm species richness shows a unimodal relationship with elevation with a peak at around 1,500 m, while the most species poor communities are located above 2,000 m. (B) Total palm abundance doesn’t have a statistically significant relationship with elevation. In both panels (A) and (B), the adjusted *R*^*2*^ and *p*-values correspond to the quadratic model (solid line), which is significantly better than the linear one (broken line). (C) A redundancy analysis (RDA) also shows that species compositional has a significant relationship with elevation, but the relationship is very weak. Gray dots represent tree plots used in the analysis (88 in total).

Do environmental conditions, particularly climate and soils, explain variation in the diversity, abundance and composition of palm communities? Regarding the effects of edaphic and climatic factors on species richness (Figure 3), we found that partitioning this variation into components (Figure 3A) shows that the shared effect was the largest (19.1%). Among the unique effects, the variation associated solely with climate was larger than that associated solely with soils (13% vs. 6.1%). However, both sets of predictors combined explained about 38.5% of the variation. In our minimum adequate regression model, we found also evidence that both soils and climate contribute to explaining variation in species richness (Figure 3C). Indeed, the best GLS model retained Clim1, Clim2, Clim3, Soils1, Soils3 and Soils 4 as significant predictors (_pseudo_ *R*^*2*^= 0.447; *P* < 0.05). The effects of climate indicate that richness increases with temperature and precipitation, but decreases with seasonality (Figure S1B, S1C and S1D). The effects of soils suggest that richness changes with concentrations of soil nutrients (Soils1), increasing toward soils with high nitrogen concentrations (Soils3). Richness also increases with cation exchange capacity (Soils4) and decreases towards acidic soils (Soils3) (Figure S2B, S2D and S2E).

**FIGURE 3.**
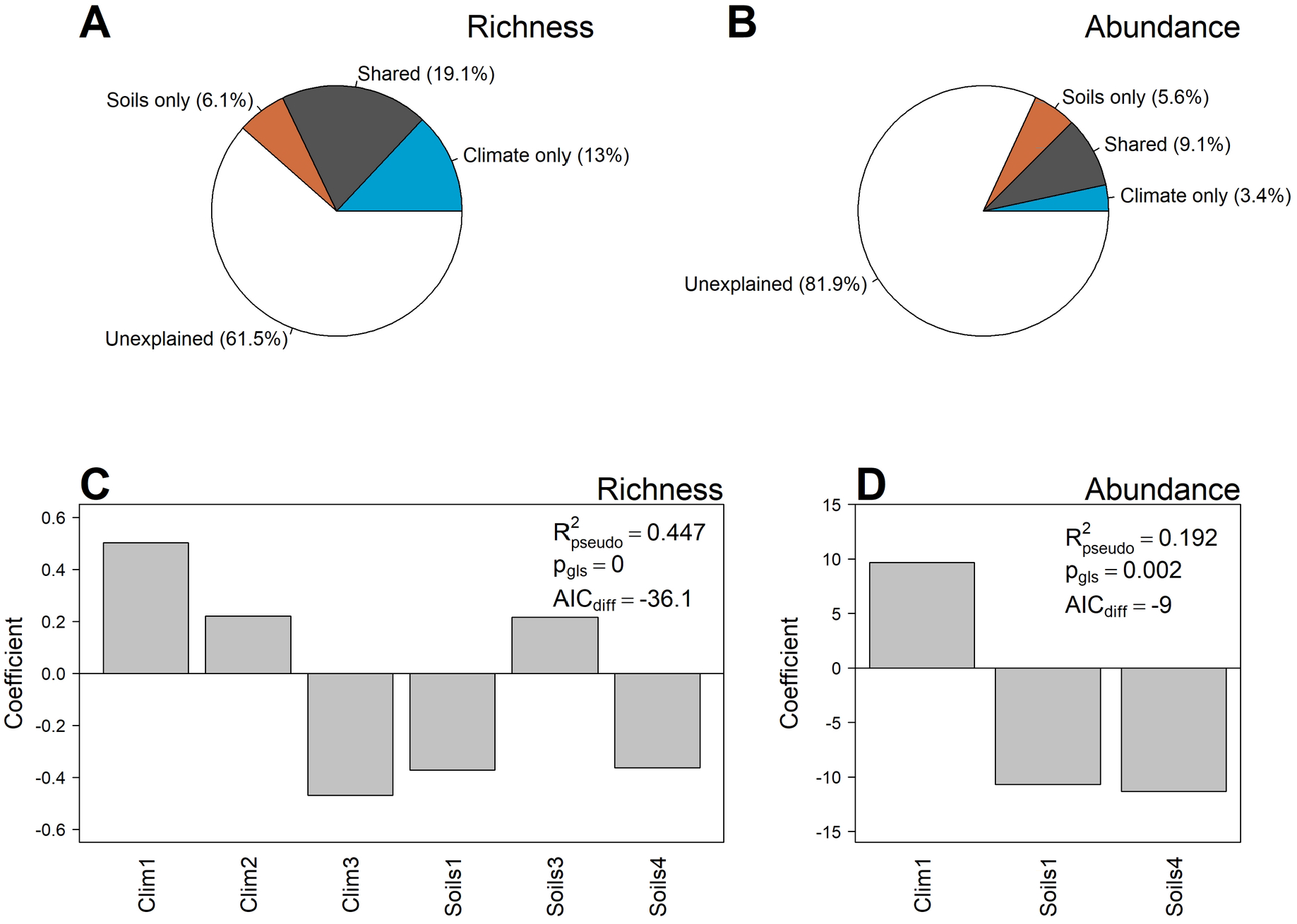
Variation in palm richness and abundance predicted by climatic and edaphic factors. Panels (A) and (B) show results of the variation partitioning analyses using all climate and soil principal components. Panels (C) and (D) show the principal components of climate and soils retained in the final models. In these panels, the pseudo-*R*^*2*^ and *p-*values from spatial GLS models are presented (A) Variation partitioning shows that less than half of the variation in richness can be explained by abiotic predictors, and that climate explains more variation than soils. (B) Variation in total palm abundance is less well explained by abiotic conditions, but both soils and climate contribute with similar fractions of variation. (C) The most important predictors for richness were climatic principal components: Clim1 and Clim3. Richness was also explained by three different soil principal components (just two are significate). (D) The most important predictors for total palm abundance included one climatic and two edaphic principal components: Clim1, Soils1 and Soils4. All of these predictors has similar relative importance.

Soils and climate explained a smaller fraction of variation in total palm abundance (Figure 3): all predictors combined accounted for only 18.1% of the variation across plots (Figure 3B). As with richness, variation partitioning showed that the shared effect of soils and climate was the largest component of variation (9.1%) while the unique effects were very small (<6%). The best GLS regression on total palm abundance retained both climatic and soil variables: Clim1, Soils1 and Soils4 (Figure 3D). The sign of these coefficients is the same as in the richness model. This suggests that total palm abundance increases with temperature (Clim1), is reduced with overall nutrient availability (Soils1), but increases with cation exchange capacity (Soils4).

When we analyzed species composition, we found that soils and climate explain about one quarter of the variation across plots (Figure 4). In contrast to the results for richness, the largest fraction of explained variation in composition is associated solely with soils (14.4%). This fraction is larger than that associated solely with climate (3.5%), or than the variation explained simultaneously by both sets of predictors (7.9%) (Figure 4A). The minimum adequate model based on redundancy analyses (RDA) retained both climatic and soil predictors (Figure 4B). Specifically, it retained all three climate principal components (Clim1 to Clim3), as well as four participant components from the soil data: Soils1, Soils2, Soils3 and Soils6.

**FIGURE 4.**
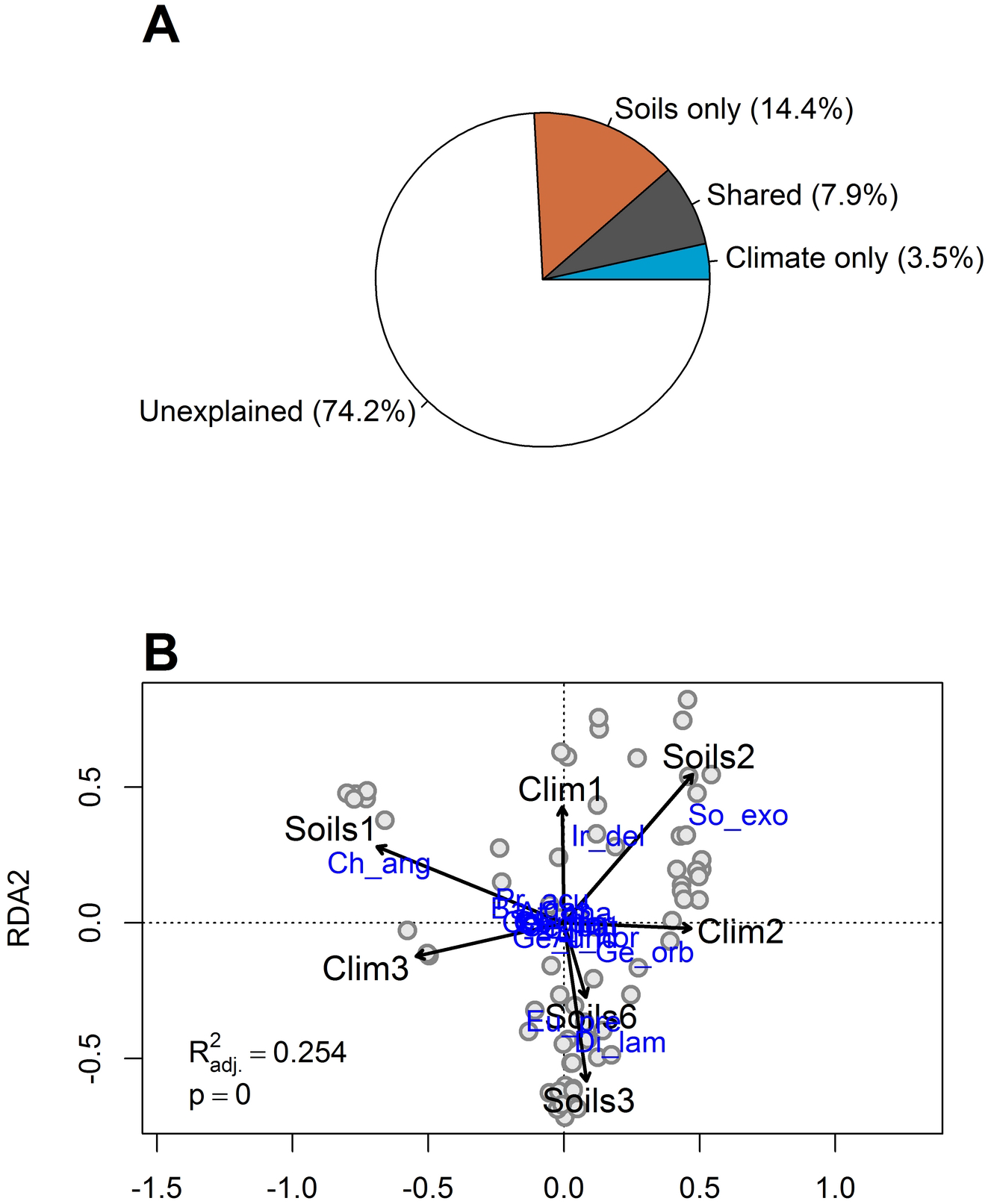
Variation in palm species composition predicted by climatic and edaphic factors. (A) Variation partitioning analysis using all principal components of climate and soils show that around one fourth of the variation can be explained. Soils explain more variation than climate. (B) An RDA with the best predictors shows the effect of both climate and soils on community composition. Four principal components of soils and three of climate were retained during variable selection (see Methods). Adjusted *R*^*2*^ and *p*-values for this RDA are also represented. Ir del = *Iriartea deltoidea*; So exo = *Socratea exorrhiza*; Ge orb = *Geonoma orbignyana*; Eu pre = *Euterpe precatoria*; Di lam = *Dictyocaryum lamarckianum*; Ch ang = *Chamaedorea angustisecta*.

Do climatic and soil conditions explain spatial variation in abundance of common species of Andean palms? We found a high degree of variability in analyses for each of the seven common species (Figure 5). Total amount of variation explained varied from nearly 60% (*C. angustisecta*) to around 10% (*G. orbignyana*). However, for two species, *A. horrida* and *G. orbignyana*, we could not find a minimum adequate model that was significantly better than a null model with only an intercept (Figure 6). Thus, we conclude that climatic and soil predictors do not account for variation in abundance of these two species. Among the remaining five species, climate seems to be more important than soils for three species (*C. angustisecta, D. lamarckianum*, and *E. precatoria*), while soils seem to be more important than climate for *I. deltoidea* and *S. exorrhiza* (Figure 5).

**FIGURE 5.**
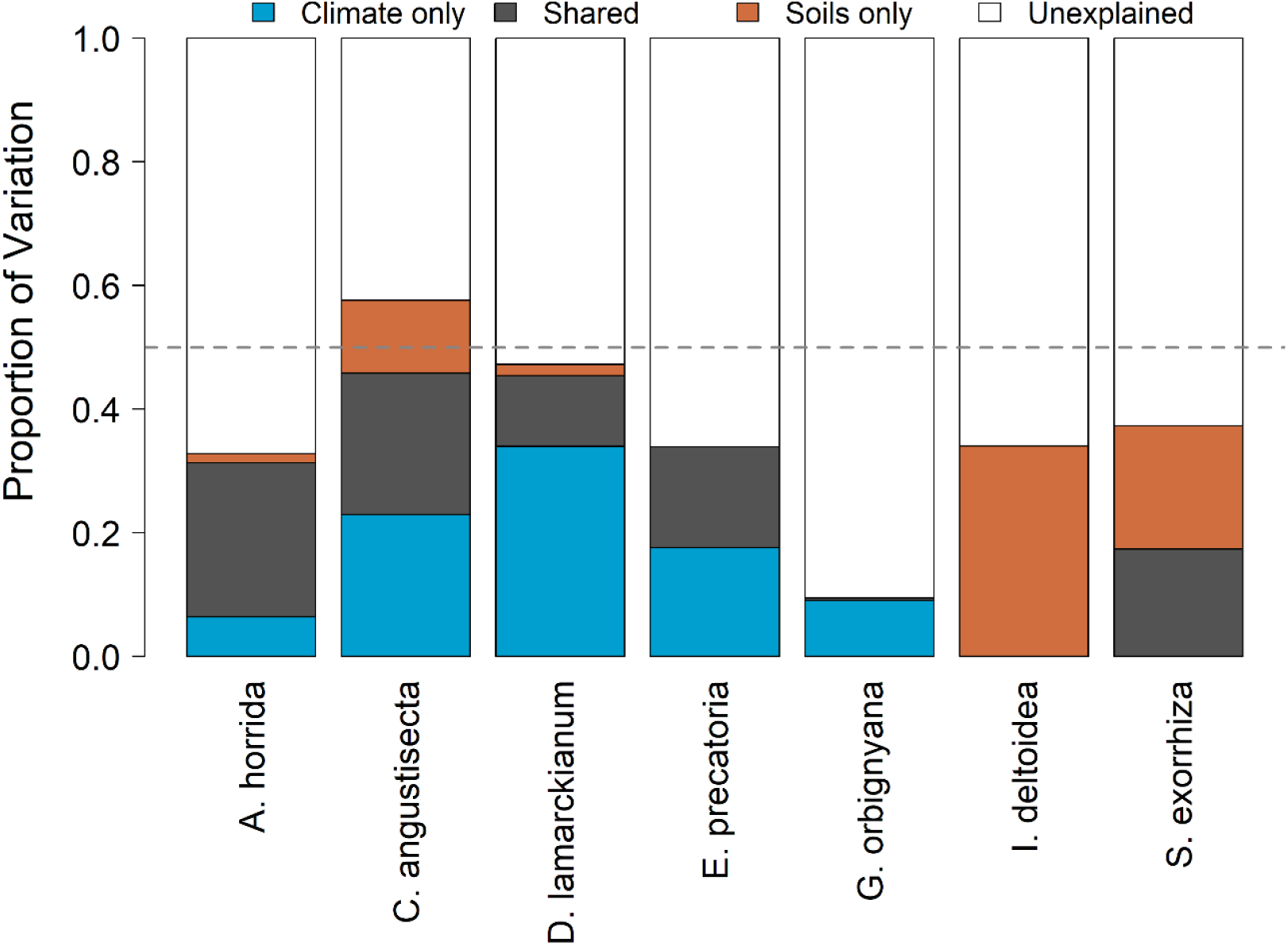
Variation partitioning between climate and soils for individual palm species. Each bar shows fractions of variation using all climate and soil predictors. Two species shows more variation explained by edaphic conditions than climate: *I. deltoidea* and *S. exorrhiza*. The other five species show more variation explained by climate than soils: *A. horrida, C. angustisecta, D. lamarckianum, E. precatoria* and *G. orbignyana*. Species also vary considerably in the total amount of variation explained. Only *C. angustisecta* have more than 50% of their variation in abundance explained.

**FIGURE 6.**
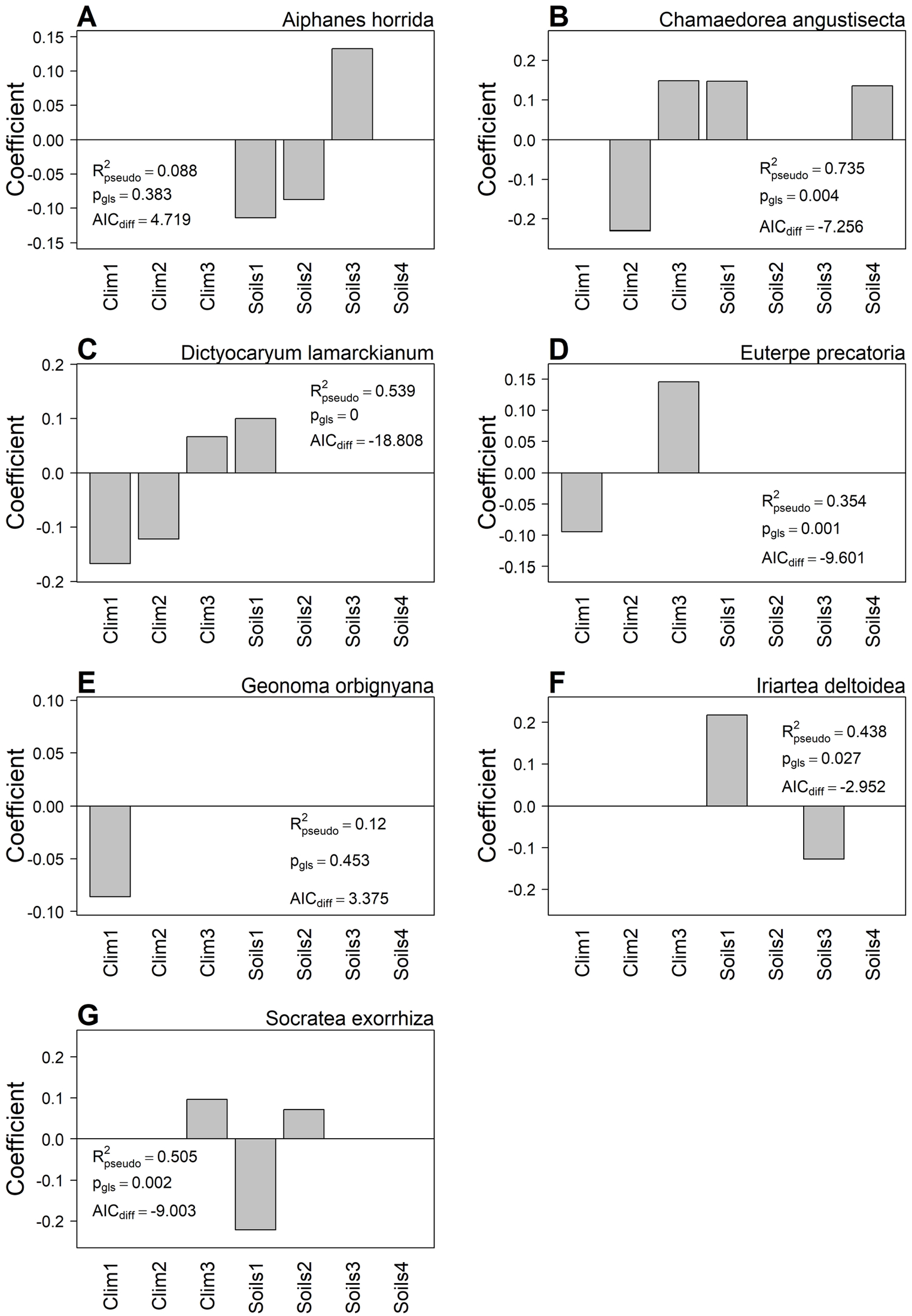
Abundance of individuals palm species predicted by climate and soil factors. Each panel shows the predictors selected in the minimum adequate model of a different common species. In each panel, the pseudo-*R*^*2*^ and *p*-values from the corresponding spatial GLS model are presented. For two species, *A. horrida* and *G. orbignyana* the GLS models are not statistically better than null models with just an intercept.

For *Chamaedorea angustisecta*, the best model (MAM) includes both two climatic and soils variables (Table 2, Figure 6B). The abundance of this species increases towards drier and more seasonal sites with soils rich in nutrients like P, Mg, Ca, K, but low in Na. The best model for *Dictyocaryum lamarckianum* includes Clim1, Clim2, Clim3 and Soils1. This species is more abundant in colder and drier sites with higher variation in temperature, and is also more abundant in sites with nutrient rich soils. The best model for *Euterpe precatoria* just contains two climate variables: Clim1 and Clim3, which means that abundance of this species is high in cooler sites with higher variation in precipitation across the year (Figure 6D). *Iriartea deltoidea* shows a significant relation with Soils1 and Soils3 (Table 2); its abundance increases towards sites with pH neutral soils and high nutrient concentration, particularly N (Figure 6F). Finally, for *Socratea exorrhiza* the best model includes Clim3, Soils1 and Soils2. This means that the abundance of this species increases in sites with high seasonality in precipitation and towards less sandy, more silty soils with low concentrations of nutrients like P, Mg, Ca and K (Figure 6G).

**Table 2.** Most important climate and soil predictors for each species. The table shows result of spatial GLS regressions with the predictors that were selected based on a search for the minimum most adequate model using AIC (see methods). The analysis was used for abundance of each species of palms that is present in fifteen or more tree-plots. For each predictor in each model, a coefficient estimate, standard error (*SE*), *t*-value and *p*-value are reported. Bold font highlights statistically significant predictors.

| Species | Variable | Estimate | *SE* | *t*-value | *p*-value |
| --- | --- | --- | --- | --- | --- |
| *Aiphanes horrida* | (Intercept) | 0.564 | 0.139 | 4.071 | **<0.001** |
|  | Soils1 | -0.114 | 0.073 | -1.559 | 0.133 |
|  | Soils2 | -0.087 | 0.049 | -1.773 | 0.090 |
|  | Soils3 | 0.133 | 0.082 | 1.625 | 0.118 |
| *Chamaedorea angustisecta* | (Intercept) | 0.692 | 0.053 | 12.962 | **<0.001** |
|  | Clim2 | -0.229 | 0.063 | -3.661 | **0.004** |
|  | Clim3 | 0.148 | 0.054 | 2.759 | **0.019** |
|  | Soils1 | 0.147 | 0.053 | 2.767 | **0.018** |
|  | Soils4 | 0.135 | 0.054 | 2.502 | **0.029** |
| *Dictyocaryum lamarckianum* | (Intercept) | 0.510 | 0.037 | 13.815 | **<0.001** |
|  | Clim1 | -0.168 | 0.036 | -4.659 | **<0.001** |
|  | Clim2 | -0.122 | 0.041 | -2.970 | **0.005** |
|  | Clim3 | 0.066 | 0.045 | 1.484 | 0.147 |
|  | Soils1 | 0.100 | 0.043 | 2.310 | **0.027** |
| *Euterpe precatoria* | (Intercept) | 0.548 | 0.043 | 12.790 | **<0.001** |
|  | Clim1 | -0.094 | 0.034 | -2.773 | **0.008** |
|  | Clim3 | 0.146 | 0.041 | 3.590 | **<0.001** |
| *Geonoma orbignyana* | (Intercept) | 0.441 | 0.052 | 8.486 | **<0.001** |
|  | Clim1 | -0.086 | 0.054 | -1.598 | 0.122 |
| *Iriartea deltoidea* | (Intercept) | 0.580 | 0.058 | 9.956 | **<0.001** |
|  | Soils1 | 0.217 | 0.064 | 3.418 | **0.004** |
|  | Soils3 | -0.127 | 0.064 | -1.999 | 0.063 |
| *Socratea exorrhiza* | (Intercept) | 0.621 | 0.039 | 15.863 | **<0.001** |
|  | Clim3 | 0.096 | 0.058 | 1.664 | 0.110 |
|  | Soils1 | -0.221 | 0.058 | -3.830 | **<0.001** |
|  | Soils2 | 0.071 | 0.042 | 1.707 | 0.101 |

## 4. DISCUSSION

### 1. Richness and composition show clear elevational gradients, but abundance does not

Our results indicate that species richness decreases non-linearly with elevation (Figure 2A), but that total palm abundance does not have a significant elevational gradient (Figure 2B). The relationship between richness and elevation is widely documented in many groups of organisms (Rahbek 1995, Lomolino 2008), including palms (Svenning 2001, Eiserhardt *et al*. 2011) and in montane forests of the Bolivian Yungas (Gerold *et al*. 2008). Despite the generality of the relationship, the shape of elevational gradients in richness is less well understood. In the literature, three types of gradients have been described: linearly decreasing, hump-shaped with an intermediate peak, or a low-elevational plateau followed by a linear decrease (Colwell & Lees 2000, McCain & Grytnes 2010). We found that the gradient is non-linear and that resembles more a low-elevation plateau. Indeed, richness seems to be somewhat constant until around 1,500 m in elevation, and then decreases rapidly to zero at about 2,500 m. We found that species composition also changes with elevation, but elevation was able to explain only 6.3% of the variation in composition. This suggests that although communities at different elevations are different, other factors beyond elevation must be more important. For example, climate, soils and other abiotic or biotic forces might be more important than simply elevation to account for changes in community composition.

We were surprised to find no clear change in total palm abundance with elevation. Neither our linear or quadratic models were significant (Figure 2B). Despite this lack of statistical support, the distribution of abundance seems nonrandom. Indeed, total palm abundance seems to peak at intermediate elevations, between 1,400 and 1,500 m. This peak of abundance at intermediate elevations may result from the dispersal of species from lower and higher elevations(Fischer *et al*. 2011). According to (Fischer *et al*. 2011), mountain forest compared to flat and lowlands the higher difference between site types per area is due to different inclinations, exposures, and geological substrates; many different micro-sites. Further studies with increased sample size and larger plots might be needed to clarify this relationship between abundance and elevation in montane palm communities (Nettesheim *et al*. 2018).

### 2. Climate explains more variation in richness, but soils explain more variation in species composition

According to our results, 13% of the variation in richness is explained by temperature (principal component Clim1) and precipitation (Clim3). This is in agreement with several studies that have shown that climate conditions have major effects on tropical palms (Baldeck *et al*. 2013, Jones *et al*. 2013, Nettesheim *et al*. 2018), and also agree more broadly with the literature that shows strong correlations between species richness and climate for many groups of organisms (Francis & Currie 2003, Currie *et al*. 2004, Tello & Stevens 2010). In our study system, climate, particularly temperature, changes dramatically between lowlands and montane regions (Jones *et al*. 2013, Nettesheim *et al*. 2018). Thus, climate might be the main driver of the elevational gradient in species richness that we observe in our data (Figure 2A). Indeed, Jones *et al*. (2011) has proposed an important role for temperature in determining the distribution of species and patterns of diversity in montane habitats. How climate shapes richness patterns is highly contentious, and many hypotheses have been proposed to explain this relationship (Hawkins *et al*. 2003, Currie *et al*. 2004, Evans *et al*. 2005). Our analyses do not attempt to disentangle these possibilities, but shows that richness patterns of an iconic group of tropical organisms respond strongly to climate across a montane region of mainly temperate environmental conditions.

While climate is the best predictor of richness patterns, edaphic conditions seem to be more strongly associated to community composition. Indeed, the fraction of variation in composition associated solely to soils is four times larger than that associated solely to climate (Figure 4A). Three principal components of soils were selected for our final model (Soils1, Soils2 and Soils3), which represent variation in a broad range of soils nutrients (mainly Soils1) and soil physical conditions (texture in Soils2 and pH in Soils3). Previous research has shown that variation in abundance of palm species depends on nutrient requirements (Svenning 2001). For example, in tropical forests across Panamá, (Turner *et al*. 2018)showed that within the same community there are species adapted to high or low phosphorous (P) concentrations. In this way, different palm species can be located at same elevation with the same climate, but in sites with contrasting soil properties (Poulsen *et al*. 2006, Blach-Overgaard *et al*. 2010). This micro-habitat differentiation could represent an ecological strategy to reduce competition among co-occurring palm species, as has been proposed for Amazonian palm communities (Eiserhardt *et al*. 2013). Our analyses for individual common species also support this view, but suggest that while community composition as a whole is mainly driven by soil properties, some individual species might respond more to climate variation.

### 3. Different common species show varying responses to climate and soil conditions

In our analyses, we found that the abundances of *I. deltoidea* and *S. exorrhiza* responded to different soil conditions. *S. exorrhiza* decreases in abundance with increasing amounts of P, Mg, Ca and K (Soils1), while it increases with sand concentration (Soils2; Figure 6). These results are in agreement with Duivenvoorden *et al*. (2005), Carlos-Copete *et al*., (2019) and Henderson *et al*. (2019), who showed that the presence of this species is within the parameters characteristic of well-drained forest soils. Our results also support findings by Cámara-Leret *et al*. (2017) who suggest that this species can grow in both rich and poor soils. *I. deltoidea* shows the opposite pattern of variation increasing in abundance with the soil characteristics represented by Soils1 (P, Mg, Ca and K), which contradict the findings of Copete *et al*. (2019). *C. angustisecta* also has some important associations with soil conditions, despite responding primarily to climate. Like *I. deltoidea, C. angustisecta* responded positively to increasing P, Mg, Ca and K, but its abundance declined with higher Na concentrations and higher cation exchange capacities.

On the other hand, we found three species, *C. angustisecta, D. lamarckianum* and *E. precatoria*, which respond mostly to climatic conditions. One pair of these species respond positively to average temperature (Clim1): *D. lamarckianum* and *E. precatoria* Both become more abundant in warmer sites. A different pair of species, *C. angustisecta* and *D. lamarckianum* increase in abundance with increasing total precipitation (Clim2). Surprisingly, all three species seem to increase in abundance with increasing seasonality in precipitation (Clim3). This effect is relatively small for *C. angustisecta* and *D. lamarckianum*, but might be important for *E. precatoria*.

Overall, our analyses on the most common palm species demonstrate a high level of idiosyncrasy in how species respond to environmental conditions. Among the seven species analyzed, two had no significant associations with climate or soils, two responded more strongly to soil conditions, and three responded mainly to climate. It is important to note that nine species were too rare to be included in these analyses, and that their environmental preferences need to be further studied.

### 4. Limitations of this study

Although we found that climate and soils explain significant fractions of variation in richness, abundance and composition of palm communities in our study region, much of the variation remains unexplained. There is a multitude of methodological and biological reasons for this. First, the resolution of the environmental data might not be the best reflection of the scales at which the environment varies and species respond to it. For example, our climate data comes from Worldclim (Fick & Hijmans 2017), which uses weather stations and satellite-derived data to create global climate surfaces. To do so, however, the data needs to be interpolated, which is difficult to do in topographically heterogeneous regions like the Andes. Similarly, we have a single measurement of soil properties per forest plot. Although our plots are small, soil conditions can change quickly at the scale of a few meters (John *et al*. 2007). Thus, our measure of soil properties might also be coarse related to how species experience these environmental conditions. Despite the deficiencies in the data, there are no better data sources available. Our study is similar to many previous studies that have tried to measure the effects of environment on species composition, so it is comparable in methodology (Myers *et al*. 2013, Arellano *et al*. 2016). Without doubt, there is need to develop better datasets to further our understanding of how species and communities respond to environmental change in remote tropical regions.

Some of the unexplained variation could also be the result of other biological processes that drive the distribution of species and assembly of communities, but that were not part of the focus of our analyses. Processes such as glaciation history, human and natural disturbance, species-species interactions, dispersal and chance could all be important. Indeed, many previous studies have highlighted one or more of these mechanisms as drivers of plant community assembly (Tuomisto, Ruokolainen, *et al*. 2003, Kraft *et al*. 2008, Antonelli *et al*. 2009, Myers *et al*. 2013). Finally, we believe that much of the unexplained variation results from simple sampling effects. This means that variation from one community to another in terms of species composition and richness is the result of the small sample sizes given by our 0.1-ha plots. These size plots are widely used because it is sometimes better to spread data-collection effort in small grain sizes (plot size) distributed across broad extents (environmental or geographic gradients). That is exactly the case in our study that covered a broad elevational gradient. This small grain size, however, introduced measurement error in describing the species composition of a site, which in turn translates into unexplained variation in statistical analyses. Despite the small size of our plots, we were able to capture clear environmental signal in the diversity and composition of palm communities in our region, which was the main objective of our study. Future work would be needed to clarify whether more intensive sampling in each community improves our ability to predict community structure using climatic and soil predictors.

## Supporting information

Supplementary results

## Data availability statement

The species and soil data used in this study are part of the Madidi Project, a long-term research collaboration aimed at studying plant diversity in the tropical Andes. The data used in our analyses correspond to version 2.2 of the Madidi Project Dataset, which is deposited in Zenodo (DOI 10.5281/zenodo.4280178). Additionally, raw data of the Madidi Project are stored and managed in Tropicos ® (https://tropicos.org/home), the botanical data database of the Missouri Botanical Garden. These data can be viewed and accessed via http://legacy.tropicos.org/Project/MDI.

## 8. ACKNOWLEDGMENTS

We thank the Herbario Nacional de Bolivia, the Dirección General de Biodiversidad, the Bolivian Park Service (SERNAP), the Madidi National Park and local communities for permits, access and collaboration in Bolivia. This project was supported by generous grants from the National Science Foundation (DEB 0101775, DEB 0743457, DEB 1836353), and the National Geographic Society (NGS 7754-04 and NGS 8047-06). Additional financial support for the Madidi Project has been provided by the Missouri Botanical Garden, the Comunidad de Madrid (Spain), the Universidad Autónima de Madrid, and the Taylor and Davidson families. This work was completed as part of the master’s thesis of the lead author and was supported by the National Science Foundation (DEB 1836353), the Missouri Botanical Garden and the Shirley Graham’s fellowship. We are grateful to Olga Martha Montiel and Shirley Graham for the support and confidence. We thank all the researchers, students and local guides that were involved in the collection of the data, particularly Carla Maldonado, Maritza Cornejo, Alejandro Araujo, Javier Quisbert and Narel Paniagua. We are thankful to all the taxonomic experts that provided identifications for plant specimens. Finally, we would like to thank Sebastián Gonzalez-Caro, Isabel Morales-Belpaire and Ramiro P. López for their valuable comments and support.

## Author Contribution Statement

FM, JST and MMR conceptualized the study. AFF, LC, AA, TM, EMM, MIL collected the data. JST and LC coordinated the Madidi Project and the data collection. AFF was responsible for the curation of herbarium specimens. FM and JST conducted the analyses and wrote the original draft. All authors contributed to revisions.

