## Supplementary results for "Relative effects of edaphic conditions and climate on palm communities in the Central Andes"

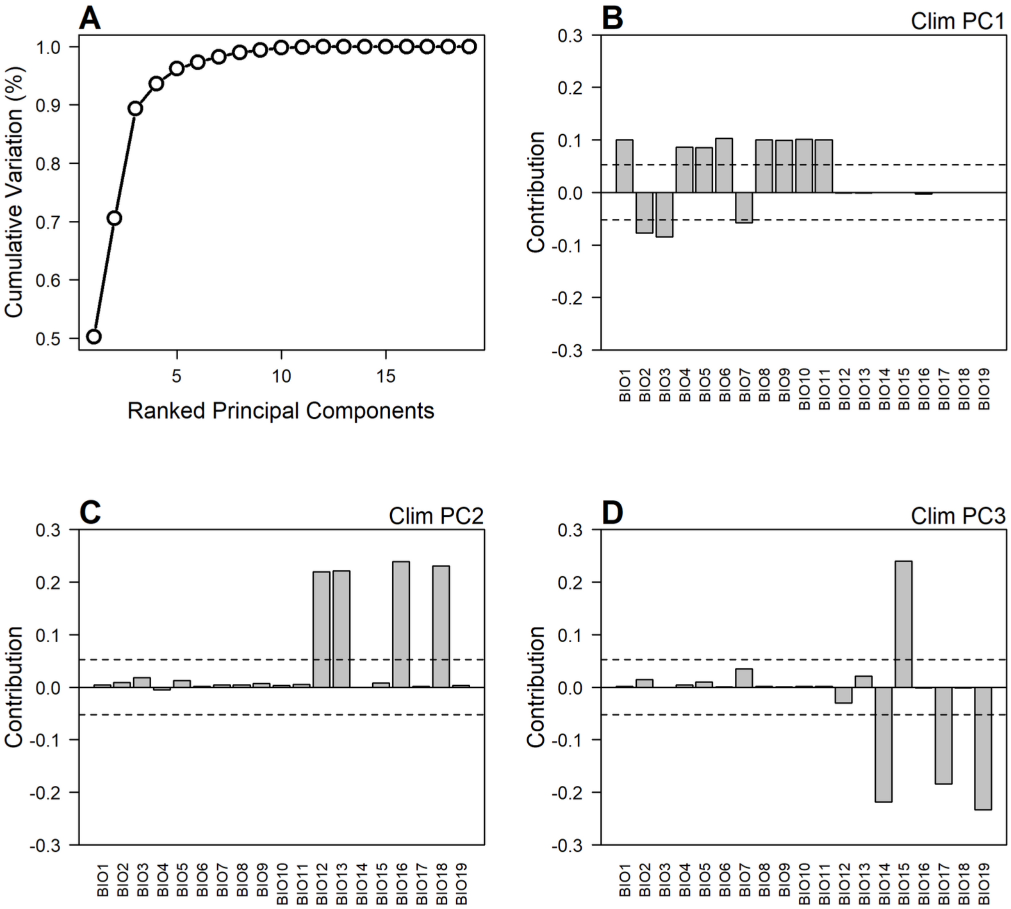


**FIGURE S1.** Summary of principal components analysis on climate data. (A) In regression analyses, we considered only the first three principal components, which account for 93% of climate variation among plots. We show loadings of each climatic variable on each of these first three principal components. All loadings can be found in Table S1. (B) The first principal component is dominated by positive loadings for most variables that represent the temperature, except mean diurnal range (BIO2), isothermality (BIO3) and temperature annual range (BIO7). (C) The second principal component mainly shows strong positive loadings for precipitation, this are related with annual precipitation (BIO12), precipitation of wettest month (BIO13), precipitation of wettest quarter (BIO16) and precipitation of warmest quarter (BIO18). (D) The third principal component is dominated by one positive loading for precipitation: precipitation seasonality (BIO15) and negative loadings: precipitation of driest month (BIO14), precipitation of driest quarter (BIO17) and precipitation of coldest quarter (BIO19).

**TABLE S1.** Loadings of principal components from climate variables in Andean forest. BIO1 = annual mean temperature; BIO2 = mean diurnal range; BIO3 = isothermality; BIO4 = temperature seasonality; BIO5 = max temperature of warmest month; BIO6 = min temperature of coldest month; BIO7 = temperature annual range; BIO8 mean temperature of wettest quarter; BIO9 = mean temperature of driest quarter; BIO10 = mean temperature of warmest quarter; BIO11 = mean temperature of coldest quarter; BIO12 = annual precipitation; BIO13 = precipitation of wettest month; BIO14 = precipitation of driest month; BIO15 = precipitation seasonality; BIO16 = precipitation of wettest quarter; BIO17 = precipitation of driest quarter; BIO18 = precipitation of warmest quarter; BIO19 = precipitation of coldest quarter. All variables come from WorldClim dataset version 2.0.

| | **PC1** | **PC2** | **PC3** | **PC4** | **PC5** | **PC6** | **PC7** | **PC8** | **PC9** | **PC10** | **PC11** | **PC12** | **PC13** | **PC14** | **PC15** | **PC16** | **PC17** | **PC18** | **PC19** |
|---|---|---|---|---|---|---|---|---|---|---|---|---|---|---|---|---|---|---|---|
| **BIO1** | 0.317 | 0.068 | 0.035 | 0.143 | 0.045 | 0.018 | 0.026 | 0.098 | 0.027 | 0.037 | 0.038 | 0.008 | 0.181 | 0.150 | 0.192 | 0.169 | 0.281 | 0.811 | 0.000 |
| **BIO2** | -0.278 | 0.093 | 0.120 | 0.461 | 0.072 | -0.026 | 0.026 | -0.027 | 0.025 | 0.137 | -0.014 | 0.041 | 0.233 | -0.091 | 0.600 | -0.481 | -0.046 | -0.054 | 0.000 |
| **BIO3** | -0.291 | 0.135 | 0.009 | 0.231 | 0.003 | 0.385 | -0.027 | 0.101 | 0.556 | 0.493 | 0.003 | 0.131 | -0.093 | 0.050 | -0.247 | 0.205 | 0.016 | 0.020 | 0.000 |
| **BIO4** | 0.293 | -0.073 | 0.067 | 0.069 | -0.240 | 0.002 | -0.246 | -0.834 | 0.178 | 0.120 | -0.136 | 0.065 | 0.070 | -0.014 | 0.020 | 0.009 | 0.090 | -0.058 | 0.000 |
| **BIO5** | 0.292 | 0.113 | 0.101 | 0.341 | 0.098 | -0.091 | 0.038 | 0.039 | -0.032 | 0.004 | -0.024 | -0.025 | -0.670 | -0.161 | 0.041 | -0.018 | 0.038 | -0.034 | 0.523 |
| **BIO6** | 0.321 | 0.044 | -0.025 | -0.054 | 0.012 | 0.082 | 0.009 | 0.108 | 0.123 | 0.070 | -0.003 | 0.012 | -0.433 | -0.148 | 0.202 | -0.143 | 0.016 | -0.041 | -0.762 |
| **BIO7** | -0.241 | 0.067 | 0.187 | 0.575 | 0.111 | -0.288 | 0.033 | -0.163 | -0.288 | -0.134 | -0.027 | -0.057 | -0.053 | 0.074 | -0.347 | 0.259 | 0.020 | 0.035 | -0.382 |
| **BIO8** | 0.317 | 0.067 | 0.035 | 0.140 | 0.040 | 0.009 | 0.020 | 0.076 | 0.027 | 0.052 | 0.030 | 0.010 | 0.299 | -0.462 | 0.028 | 0.323 | -0.673 | 0.005 | 0.000 |
| **BIO9** | 0.314 | 0.084 | 0.032 | 0.142 | 0.076 | 0.021 | 0.049 | 0.179 | 0.023 | 0.038 | 0.042 | -0.009 | 0.367 | -0.396 | -0.396 | -0.274 | 0.505 | -0.221 | 0.000 |
| **BIO10** | 0.318 | 0.055 | 0.037 | 0.129 | 0.030 | 0.026 | 0.016 | 0.052 | 0.036 | 0.033 | 0.023 | 0.038 | 0.054 | 0.550 | -0.354 | -0.512 | -0.416 | 0.055 | 0.000 |
| **BIO11** | 0.316 | 0.070 | 0.034 | 0.148 | 0.059 | 0.019 | 0.047 | 0.160 | 0.009 | 0.020 | 0.046 | 0.018 | 0.172 | 0.488 | 0.310 | 0.410 | 0.163 | -0.529 | 0.000 |
| **BIO12** | -0.036 | 0.468 | -0.174 | -0.029 | 0.060 | -0.132 | -0.220 | -0.008 | 0.316 | -0.189 | -0.176 | -0.713 | 0.045 | 0.031 | 0.015 | -0.011 | -0.018 | -0.003 | 0.000 |
| **BIO13** | -0.032 | 0.471 | 0.144 | -0.118 | -0.134 | 0.225 | 0.136 | -0.221 | -0.231 | 0.070 | 0.731 | -0.115 | -0.036 | -0.008 | 0.006 | -0.009 | 0.001 | -0.006 | 0.000 |
| **BIO14** | 0.008 | -0.005 | -0.468 | 0.025 | 0.544 | 0.077 | 0.536 | -0.337 | 0.177 | -0.182 | 0.063 | 0.091 | 0.011 | -0.001 | 0.005 | 0.001 | -0.001 | 0.000 | 0.000 |
| **BIO15** | -0.010 | 0.092 | 0.490 | -0.046 | 0.086 | 0.635 | 0.163 | -0.060 | -0.077 | -0.387 | -0.387 | -0.052 | 0.013 | -0.010 | 0.007 | 0.003 | -0.005 | 0.003 | 0.000 |
| **BIO16** | -0.051 | 0.489 | -0.038 | -0.030 | -0.148 | -0.189 | -0.152 | 0.060 | 0.240 | -0.459 | -0.017 | 0.634 | 0.006 | -0.017 | 0.000 | 0.004 | 0.007 | 0.003 | 0.000 |
| **BIO17** | 0.013 | 0.036 | -0.429 | 0.257 | -0.708 | 0.167 | 0.395 | 0.061 | -0.119 | -0.079 | -0.177 | -0.084 | -0.007 | -0.006 | -0.007 | 0.000 | 0.001 | 0.002 | 0.000 |
| **BIO18** | 0.008 | 0.481 | -0.028 | -0.257 | 0.115 | -0.149 | 0.180 | -0.066 | -0.366 | 0.503 | -0.472 | 0.138 | 0.020 | 0.020 | -0.012 | 0.013 | 0.005 | 0.003 | 0.000 |
| **BIO19** | 0.008 | 0.059 | -0.483 | 0.170 | 0.199 | 0.433 | -0.581 | 0.007 | -0.399 | -0.032 | -0.011 | 0.098 | -0.017 | -0.009 | 0.002 | -0.001 | 0.002 | -0.001 | 0.000 |

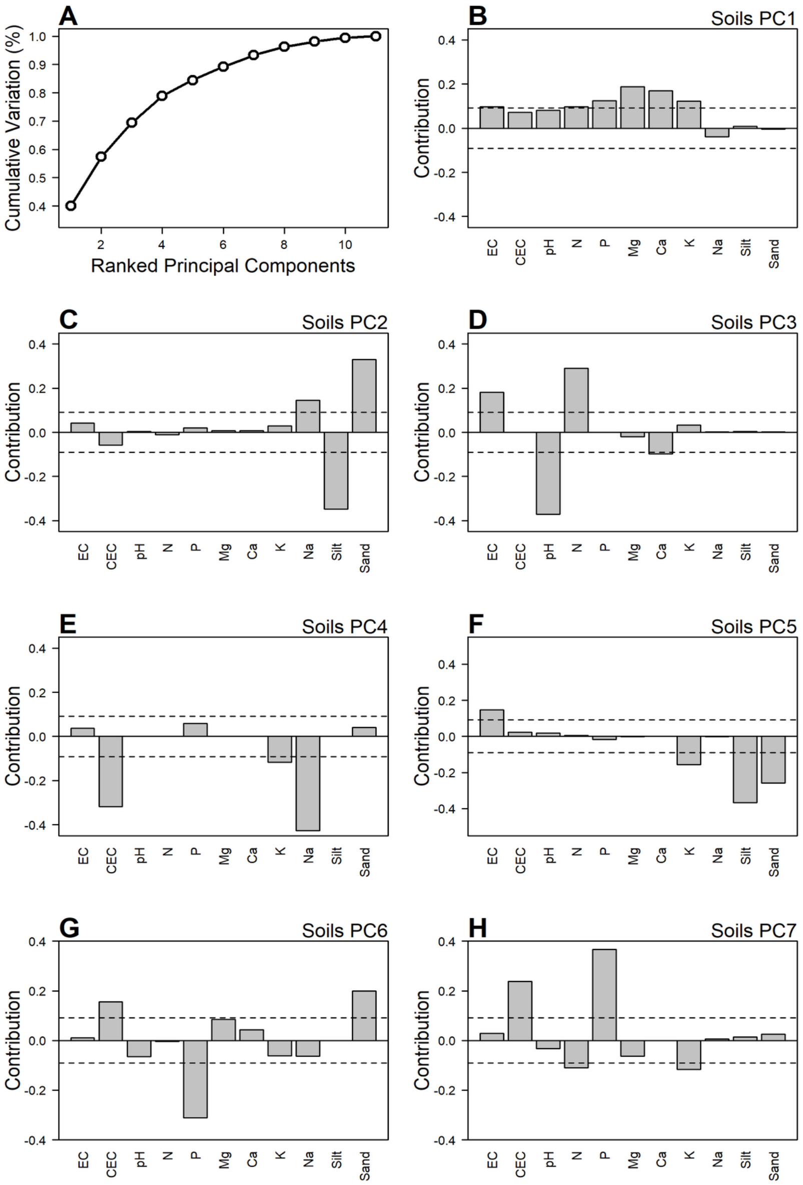


**FIGURE S2.** Summary of principal components analysis on soils data. (A) In regression analyses, we considered the first seven principal components, which account for 92% of edaphic variation among plots. All loadings can be found in Table S2. (B) The first principal component is dominated by positive loadings for most variables, the strongest loadings are P, Mg, Ca and K. (C) The second principal component shows positive and strongest loadings for Na and sand, the principal negative loading is silt. (D) The third principal component shows two strongest positive loadings: electro conductivity and N, and the negative loadings is pH. (E) The fourth principal component shows just strong negative loadings, these are related with CEC, K and Na. (F) The fifth principal component is dominated by negative loadings, the strongest positive loading is electro conductivity and negative loadings are related with K, silt and sand. (G) The sixth principal component shows a strongest positive loading for cationic exchangeable capacity and Sand, and the strongest negative loadings is phosphorus. (H) The seventh principal component shows two strongest positive loadings: cationic exchangeable capacity and phosphorus and negative loadings are related with N and K.

**TABLE S2.** Loadings of principal components from soils variables in Andean forests. Positive and strongest loadings are indicated in red, negative in blue. Just for the first seven principal components (PC). EC = Electro conductivity; CIC = Cationic interchange capacity; pH = potential of hydrogen; C = Carbon; N = Nitrogen; P = Phosphorus; Mg = Magnesium; Ca = Calcium; K = Potasium: Na = Sodium; Sil = Silt; San = Sand.

| | **PC1** | **PC2** | **PC3** | **PC4** | **PC5** | **PC6** | **PC7** | **PC8** | **PC9** | **PC10** | **PC11** |
|---|---|---|---|---|---|---|---|---|---|---|---|
| **EC** | **0.311** | 0.206 | **0.426** | 0.191 | **0.385** | 0.104 | 0.171 | 0.454 | 0.186 | 0.458 | 0.065 |
| **CEC** | 0.268 | -0.240 | -0.020 | **-0.564** | 0.152 | **0.395** | **0.488** | -0.334 | -0.069 | 0.139 | 0.011 |
| **pH** | 0.284 | 0.053 | **-0.610** | 0.027 | 0.140 | -0.255 | -0.182 | -0.247 | 0.447 | 0.397 | 0.095 |
| **N** | **0.311** | -0.103 | **0.539** | -0.013 | 0.080 | -0.060 | **-0.331** | -0.430 | 0.428 | -0.323 | -0.103 |
| **P** | **0.352** | 0.141 | 0.026 | 0.240 | -0.134 | **-0.558** | **0.606** | -0.134 | -0.127 | -0.242 | 0.094 |
| **Mg** | **0.434** | 0.084 | -0.140 | 0.017 | -0.048 | 0.293 | -0.252 | 0.153 | -0.181 | -0.316 | 0.691 |
| **Ca** | **0.412** | 0.084 | **-0.314** | -0.006 | 0.000 | 0.208 | -0.001 | 0.357 | 0.075 | -0.362 | -0.646 |
| **K** | **0.350** | 0.169 | 0.180 | **-0.343** | **-0.396** | -0.248 | **-0.341** | -0.001 | -0.432 | 0.380 | -0.182 |
| **Na** | -0.195 | **0.382** | 0.034 | **-0.654** | -0.045 | -0.252 | 0.078 | 0.304 | 0.388 | -0.215 | 0.167 |
| **Silt** | 0.093 | **-0.590** | 0.054 | 0.034 | **-0.605** | 0.003 | 0.116 | 0.317 | 0.364 | 0.124 | 0.105 |
| **Sand** | -0.059 | **0.575** | 0.033 | 0.200 | **-0.508** | **0.447** | 0.157 | -0.274 | 0.230 | 0.115 | -0.026 |

**TABLE S3.** Interpretation of the soils principal component axes. For each axis, the variables with significant positive or negative loads are indicated and used as the basis for the interpretation of each axis.

| PC | Positive variables | Negative variables | Interpretation |
|---|---|---|---|
| **Soils1** | Phosphorus<br>Magnesium<br>Calcium<br>Potassium | | A gradient from soils with low concentration in P, Mg, Ca and K to soils with a higher concentration |
| **Soils2** | Sodium<br>Sand | Silt | A gradient from soils with high percentage of silt to soils with high concentration of Na and sand |
| **Soils3** | Electro conductivity<br>Nitrogen | pH | Gradient from acidic soils to soils with high electro conductivity and high concentration of N |
| **Soils4** | | Cationic exchangeable capacity<br>Potassium<br>Sodium | A gradient from soils with high cationic exchangeable capacity, high concentrations of K and Na to soils with lowest concentrations |
| **Soils5** | Electro conductivity | Potassium<br>Silt<br>Sand | A gradient from soils with high percentage of silt and sand and high concentration of K to soils with high electro conductivity |
| **Soils6** | Cationic exchangeable capacity<br>Sand | Phosphorus | A gradient from soils with high concentration of P to soils with high cationic exchangeable capacity and high percentage of sand |
| **Soils7** | Cationic exchangeable capacity<br>Phosphorus | Nitrogen<br>Potassium | A gradient from soils with high concentration of N and K to soils with high concentration of P and high cationic exchangeable capacity |

**TEXT S1. Brief description of effect of soils conditions on plant distribution. –** The availability of soil nutrients limits plant species distribution because it provides input to tissue formation and metabolic activities (Bray & Kurtz 1945, Ra*ven et* al. 2005). Soil is the primary nutrient medium for plants. The main nutrients obtained from soils are nitrogen (N) and phosphorus (P) that are basic components of proteins, nucleotides and enzymes (Steven*son et* al. 1982, Ra*ven et* al. 2005). A deficit of N produces metabolic failures in metabolic machinery and can cause leaf death. Some plants are evolved strategies to acquire N in low nutrient soils such as symbiosis with nitrogen fixing microorganisms (e.g. *Rhizobium* in Fabaceae trees) (Steven*son et* al. 1982, Ra*ven et* al. 2005). P is integral part of the adenosine triphosphate (ATP) molecule. Its deficit reduces metabolism and protein formation. Thus, P has been identified as a limiting factor for tropical tree growth. However, P limitation is not pervasive for all tropical species because many of them are adapted to low concentrations of this nutrient (Bray & Kurtz 1945, Ra*ven et* al. 2005).

Regarding other macronutrients; magnesium (Mg) is a component of the chlorophyll molecule, activator of many enzymes (Van R*aij et* al. 1986, Ra*ven et* al. 2005). Mg deficit produces mottled or chlorotic leaves, sometimes with necrotic spots, leaf tips and margins turned upward, and slender stems. Calcium (Ca) is a component of cell walls, enzyme cofactors, involved in cellular membrane permeability, component of calmodulin, a regulator of membrane and enzyme activities among other important molecules (Graust*ein et* al. 1977, Van R*aij et* al. 1986, Ra*ven et* al. 2005). A deficit on Ca causes shoot and root tips to die, hooked young leaves that eventually die back at tips and margins. Potassium (K) is involved in osmosis and ionic balance and stomata dynamics, and it is also an activator of many enzymes. A deficit in K causes mottled or chlorotic leaves with small spots of necrotic tissue at tips and margins, weak, narrow stems (Conti 1992, Ra*ven et* al. 2005).

In the other hand, the weakly bound cations can be replaced by other cations and thus released into the soil solution, where they become available for the plant growth. This process is called cation exchange (Ma & Eggleton 1999). pH is a measure of the acidity and alkalinity in soils. pH levels range from 0 to 14, with 7 being neutral, below 7 acidic and above 7 alkaline (Corwin & Lesch 2003, Raven *et al.* 2005). The acidity or alkalinity of soil is related to the availability of inorganic nutrients for plant growth (Ma & Eggleton 1999). Soils vary widely in pH, and many plants have a narrow range of tolerance on this scale. In alkaline soils, some cations are precipitated, and such elements as iron, manganese, copper, and zinc may thereby become unavailable to plants (Ma & Eggleton 1999, Meier 2006). Electro-conductivity (EC), measures the capacity of the soil to conduct electricity by salts. Therefore, the EC measures the concentration of soluble salts present in the soil (Corwin & Lesch 2003, Sudd*uth et* al. 2005). High salinity facilitates electric interchange and nutrient mobilization (Corwin & Lesch 2003). An excess of sodium (Na) in the soils has adverse effects on the growth of plants and soil structure (Sher*aed et* al. 1976, Ra*ven et* al. 2005) by reducing oxygen availability and oxygenation capacity on the roots (Sher*aed et* al. 1976). At high Na concentrations, the soil becomes hard, crusty, which can restricts germination and the normal growth of roots (Sher*aed et* al. 1976). Moreover, sodic soils are susceptible to erosion, which generates loss of soil and nutrients (Sher*aed et* al. 1976).

859–870.

Van Raij, B., J. A. Quaggio, and N. M. Da Silva. 1986. Extraction of phosphorus, potassium, calcium, and magnesium from soils by an ion-exchange resin procedure. Commun. Soil Sci. Plant Anal. 17: 547–566.

Raven, P. H., R. F. Evert, S. E. Eichhorn, and others. 2005. Biology of plants. Macmillan.

Schlindwein, G., A. Tonietto, A. D. Abichequer, A. C. de Azambuja, B. B. Lisboa, and L. K. Vargas. 2017. Pindo Palm fruit yield and its relationship with edaphic factors in natural populations in Rio Grande do Sul. Ciência Rural 47: 1–7.

Sesnie, S. E., B. Finegan, P. E. Gessler, Z. Ramos, and C. Rica. 2009. Landscape-Scale Environmental and Floristic Variation in Costa Rican Old-Growth Rain Forest Remnants. Biotropica 41: 16–26.

Sheraed, J. L., L. P. Dunnigan, and R. S. Decker. 1976. Identification and nature of dispersive soils. J. Geotech. Geoenvironmental Eng. 102.

Smith, T. W., and J. T. Lundholm. 2010. Variation partitioning as a tool to distinguish between niche and neutral processes. Ecography (Cop.). 33: 648–655.

Stevenson, F. J., J. M. Bremner, and others. 1982. Nitrogen in agricultural soils. American Society of Agronomy Madison-Wisconsin Wisconsin.

Sudduth, K. A., N. R. Kitchen, W. J. Wiebold, W. D. Batchelor, G. A. Bollero, D. G. Bullock, D. E. Clay, H. L. Palm, F. J. Pierce, R. T. Schuler, and K. D. Thelen. 2005. Relating apparent electrical conductivity to soil properties across the north-central USA. Comput. Electron. Agric. 46: 263–283.

Svenning, J.-C. 2001. On the role of microenvironmental heterogeneity in the ecology and diversification of Neotropical rain forest palms (Arecaceae). Bot. Rev. 67: 1–53.

Tuomisto, H., A. D. Poulsen, K. Ruokolainen, R. C. Moran, C. Quintana, J. Celi, and G. Cañas. 2003. Linking floristic patterns with soil heterogeneity and satellite imagery in Ecuadorian Amazonia. Ecol. Appl. 13: 352–371.

Turner, B. L., T. Brenes-Arguedas, and R. Condit. 2018. Pervasive phosphorus limitation of tree species but not communities in tropical forests. Nature 555: 367–370. Available at: http://dx.doi.org/10.1038/nature25789.

Venables, W. N., and B. D. Ripley. 2002. Modern applied statistics (Fourth S., editor) New York.

Wright, S. J. 2019. Plant responses to nutrient addition experiments conducted in tropical forests Available at: https://onlinelibrary.wiley.com/doi/abs/10.1002/ecm.1382.
